# Prioritized polycystic kidney disease drug targets and repurposing candidates from pre-cystic and cystic mouse *Pkd2* model gene expression reversion

**DOI:** 10.1101/2022.12.02.518863

**Authors:** Elizabeth J. Wilk, Timothy C. Howton, Jennifer L. Fisher, Vishal H. Oza, Ryan T. Brownlee, Kasi C. McPherson, Hannah L. Cleary, Bradley K. Yoder, James F. George, Michal Mrug, Brittany N. Lasseigne

## Abstract

Autosomal dominant polycystic kidney disease (ADPKD) is one of the most prevalent monogenic human diseases. It is mostly caused by pathogenic variants in *PKD1* or *PKD2* genes that encode interacting transmembrane proteins polycystin-1 (PC1) and polycystin-2 (PC2). Among many pathogenic processes described in ADPKD, those associated with cAMP signaling, inflammation, and metabolic reprogramming appear to regulate the disease manifestations. Tolvaptan, a vasopressin receptor-2 antagonist that regulates cAMP pathway, is the only FDA-approved ADPKD therapeutic. Tolvaptan reduces renal cyst growth and kidney function loss, but it is not tolerated by many patients and is associated with idiosyncratic liver toxicity. Therefore, additional therapeutic options for ADPKD treatment are needed. As drug repurposing of FDA-approved drug candidates can significantly decrease the time and cost associated with traditional drug discovery, we used the computational approach signature reversion to detect inversely related drug response gene expression signatures from the Library of Integrated Network-Based Cellular Signatures (LINCS) database and identified compounds predicted to reverse disease-associated transcriptomic signatures in three publicly available *Pkd2* kidney transcriptomic data sets of mouse ADPKD models. We focused on a pre-cystic model for signature reversion, as it was less impacted by confounding secondary disease mechanisms in ADPKD, and then compared the resulting candidates’ target differential expression in the two cystic mouse models. We further prioritized these drug candidates based on their known mechanism of action, FDA status, targets, and by functional enrichment analysis. With this in-silico approach, we prioritized 29 unique drug targets differentially expressed in *Pkd2* ADPKD cystic models and 16 prioritized drug repurposing candidates that target them, including bromocriptine and mirtazapine, which can be further tested in-vitro and in-vivo. Collectively, these indicate drug targets and repurposing candidates that may effectively treat pre-cystic as well as cystic ADPKD.

Graphical abstract of the study created with Biorender.com.

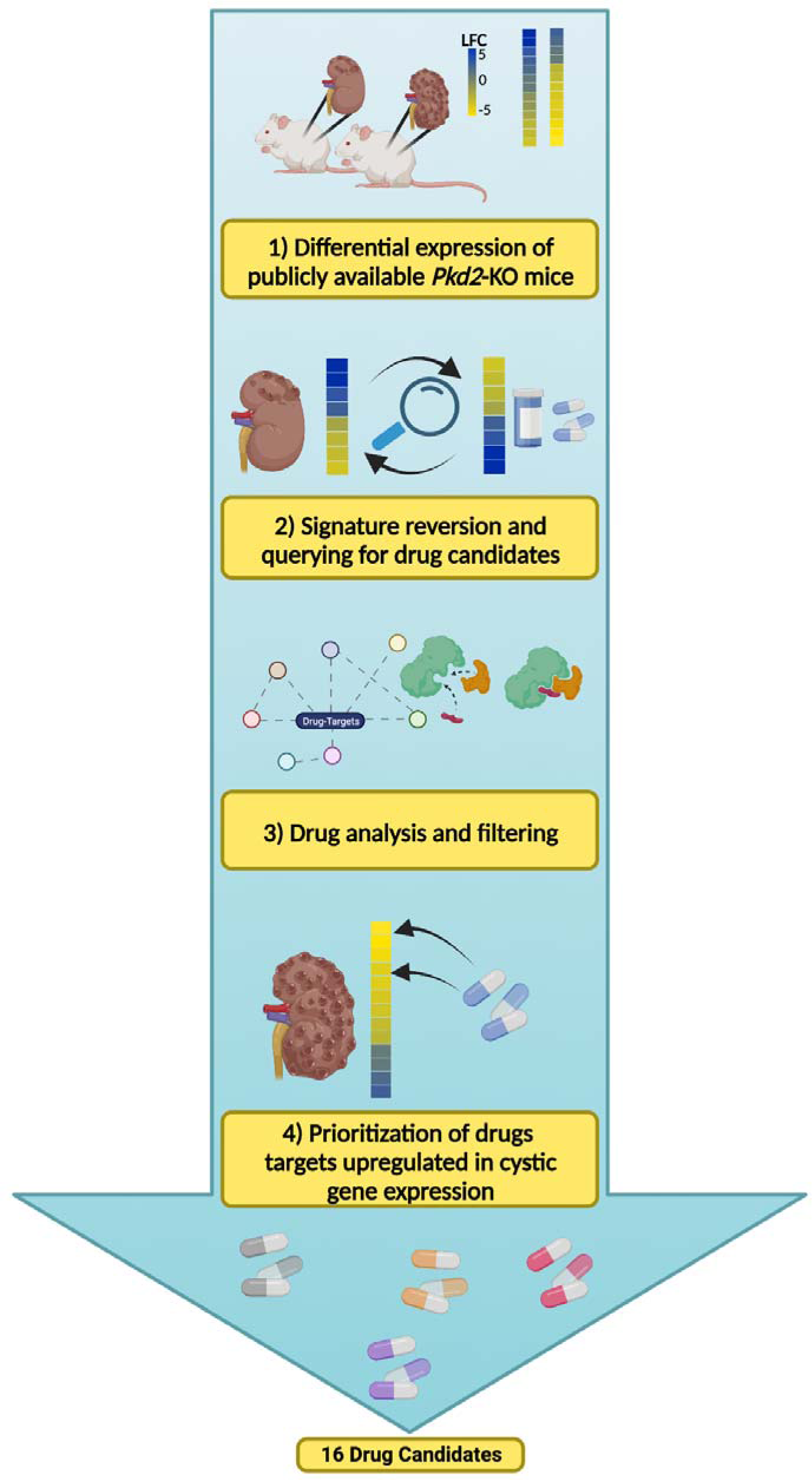

## Background

Autosomal dominant polycystic kidney disease (ADPKD) is a potentially life-threatening disorder with renal and extrarenal manifestations. It is one of the most prevalent monogenic human diseases, impacting approximately 1 in 400 to 1 in 1,000 individuals.(Cordido et al., 2017) In addition to end-stage renal disease (ESRD), comorbidities include hypertension, extrarenal cysts, and intracranial aneurysms and hemorrhaging.(Cordido et al., 2017) ADPKD is primarily caused by variants in either the *PKD1* or *PKD2* gene which encode the transmembrane proteins polycystin-1 (PC1) and polycystin-2 (PC2), respectively.(Cordido et al., 2017) While many molecular processes have been associated with ADPKD, the interrelated processes of cyst expansion, inflammation, and metabolic reprogramming have all been identified as key molecular processes for regulating disease manifestation.(Cordido et al., 2017; Podrini et al., 2020) Variants in *PKD1* and *PKD2* disrupt extracellular sensing and calcium transport by the protein products, leading to decreased intracellular calcium and triggering upregulation of cyclic adenosine monophosphate (cAMP).(Lemos & Ehrlich, 2018)(Chebib et al., 2015) This increased cAMP leads to increased fluid secretion and proliferation, resulting in fluid accumulation in the cyst lumen. Inflammation in ADPKD is initiated early in disease pathogenesis, as evidenced by cytokine and other proinflammatory signaling molecules in patients’ urine, likely caused by cellular alterations from polycystin dysfunction.(Karihaloo, 2016) Additionally, inflammation plays a role in disease progression through macrophage infiltration. Macrophage accumulation has been linked to cyst expansion, and macrophage depletion in mice causes decreased cyst growth, improved renal function, and reduced cell proliferation in cyst-lining cells.(Karihaloo, 2016) PC1 and PC2 interact directly and indirectly with mitochondria, resulting in a “Warburg-like” aerobic glycolysis effect in ADPKD as well as dysregulation in the pentose phosphate pathway, fatty acid biosynthesis, oxidative phosphorylation, and other metabolic pathways.(Podrini et al., 2020; Seeger-Nukpezah et al., 2015) Tolvaptan, the only current FDA-approved drug for ADPKD, is a vasopressin receptor-2 antagonist associated with cAMP regulation.(Vasileva et al., 2021) Vasopressin receptor antagonists (vaptans) have been the primary therapeutic focus to treat ADPKD.(Vasileva et al., 2021) While tolvaptan reduces cyst growth and formation, slowing disease progression, it is expensive and treatment is associated with adverse events including polyuria, polydipsia, elevated liver enzymes, and liver toxicity leading to non-compliance and withdrawal from treatment.(Bellos, 2021; Pellegrino et al., 2019) Therefore, additional therapeutic options are needed for ADPKD treatment.

Drug repurposing of FDA-approved drug candidates can significantly decrease the time and cost associated with traditional drug discovery by bypassing phase I and II clinical trials (the estimated average in the US is $300 million compared to ∼$2-3 billion for a new drug).(Pushpakom et al., 2019) Here, we use the computational approach signature reversion to identify compounds capable of reversing disease-associated transcriptomic signatures to prioritize drug repurposing candidates. This approach of identifying drug repurposing candidates via signature reversion has been previously successfully applied and translated to preclinical models across multiple diseases including pulmonary arterial hypertension, inflammatory bowel disease, hepatocellular carcinoma, neutrophilic bronchial asthma, skeletal muscle atrophy, and dyslipidemia.(Dudley et al., 2011; Gu et al., 2021; Kunkel et al., 2011; Regan-Fendt et al., 2020; Shin et al., 2015; Wagner et al., 2015). As gene expression profiles from ADPKD patients and preclinical models have significantly altered transcriptomes, our approach presents an opportunity to compare disease signatures to drug response signatures from treated cell lines and identify drug candidates that may reverse ADPKD-associated cellular phenotypes, ultimately slowing or reducing kidney cyst growth. Additionally, drug selection based on genetic evidence has become more common and is effective. For example, approximately 66% of FDA-approved drugs in 2021 were supported by demonstrated associations between genetic phenotype and drug targets.(Ochoa et al., 2022) This further emphasizes the utility of omics profiles to identify the right drug for the right patient at the right time.

Here, our focus was on *Pkd2*, due to public high-quality data set availability and similarity to *Pkd1* in leading to an ADPKD-like phenotype in preclinical mouse models. From those data sets, we prioritized drug repurposing candidates for ADPKD by detecting inversely related drug response gene expression signatures from the Library of Integrated Network-Based Cellular Signatures (LINCS) database compared to pre-cystic kidney gene expression. Drug candidates were further prioritized based on targets upregulated in cystic kidney gene expression as well as their known mechanism of action (MOA), FDA status, targets, side effects, and by functional enrichment analysis.

## Methods

### Data Acquisition and Processing

Three previously published kidney C57BL6 mice RNA-Seq data sets that contained either *Pkd2^fl^*^/fl^;*Pax8*rtTA;TetO-Cre (GSE149739(Zhang et al., 2021)) or *Pkhd1*-cre;*Pkd2*^F/F^ (GSE134719(Lee et al., 2019) and GSE69556(Lakhia et al., 2016)) (Additional file 1) with matched controls and sexes were downloaded from SRA using SRA-Toolkit version 2.10.7. We selected these data sets because they each were derived from *Pkd2* knock-out kidney mouse samples on C57BL6 backgrounds and had matched controls. GSE149739 samples (P70) were from mice induced at postnatal day 28 for 2 weeks and euthanized at postnatal day 70, when, the authors had noted, there was mild tubule dilation but prior to overt cyst development (i.e., pre-cystic). For the P70 data set, 3 *Pkd2* knock-out (KO) samples and 3 matched wild-type samples were used in our study. *Pkhd1*/cre;*Pkd2*^F/F^ mice and are under tissue-specific *Pkhd1*-promoter regulation using the previously described *Pkhd1*/cre strain, which results in rapidly develop renal cysts due to constitutive knockout of *Pkd2*.(Williams et al., 2014) As a result, GSE134719 (P28) and GSE69556 (P21) kidneys were collected at postnatal day 28 and 21, respectively. For the analyses here, we used 11 *Pkd2* KO and 12 wild-type samples for the P28 data set, and 3 KO and 4 wild-type samples for the P21 data set.

In short, we processed FASTQ files for all samples on the UAB Cheaha supercomputer with the nf-core/rnaseq(Ewels et al., 2019, 2020) pipeline version 3.6 using the STAR/Salmon route for alignment, adapter trimming, quality control (QC), and gene count calculations. We aligned reads to the GENCODE mouse reference genome (mm10 release M24). All other analyses were performed on a 2019 MacBook Pro (2.4 GHz Intel Core i9, 64 GB DDR memory, 2TB SSD storage) using R version 4.2.0 RStudio 2021.09.0+351 "Ghost Orchid" Release (2021-09-20) for macOS.

### Differential Expression Analysis

For each of the three datasets, we performed differential expression analysis using DESeq2 (version 1.34.0)(Love et al., 2014) on the STAR/Salmon merged gene counts from the nf-core pipeline outputs. Differentially expressed genes were selected based on a threshold of an absolute log2 fold change (LFC) greater than 1.1 (for pre-cystic P70 data set) or 2.0 (for cystic P21 and P28 data sets) and an adjusted p-value less than 0.05 after apeglm LFC shrinkage.(Zhu et al., 2019) The LFC cutoff varied by data set to ensure a signature of 100-200 genes, which Yang et al. previously found to be the optimal gene signature size for signature reversion.(Yang et al., 2022) While a LFC threshold > 2 allowed for a total signature of 100-200 genes (126 genes for P21 and 189 genes for P28) for the cystic data sets; there were far fewer genes passing this stringent cutoff with the pre-cystic data set. The study by Yang et al. benchmarked transcriptomic signature compositions for drug repurposing and found the optimal signature to be dependent more on size (100-200 genes) than stringent LFC.(Yang et al., 2022) With this in mind, we lowered the pre-cystic LFC cutoff to > 1.1 and < -1.1 compared to cystic data sets, resulting in a signature of 130 genes for the P70 pre-cystic data set. We used BiomaRt to annotate gene descriptions for the differentially expressed genes and convert from Ensembl to Entrez and HGNC symbols (version 2.50.3).(Durinck et al., 2005, 2009) The top differentially expressed genes by LFC were used to select rlog normalized counts and plotted as heatmaps using complete linkage hierarchical clustering.

### Functional Enrichment Analysis (FEA)

We then analyzed Mouse Ensembl genes (C57BL_6NJ_v1) that met the differentially expressed LFC cutoff for pathway and functional enrichment, with upregulated and downregulated genes analyzed separately. We used the R package gprofiler2 (version 0.2.1)(Raudvere et al., 2019; Reimand et al., 2007) to identify significantly enriched pathways using the Gene Ontology (GO), Reactome, and WikiPathways gene set sources (term sizes filtered for > 5 and < 1,000). We applied the Bonferroni Procedure for multiple hypothesis correction and used a p-adjusted threshold of 0.05. The background gene list included all measured genes before filtering. For plotting, we used enrichment terms prioritized by recall value, the ratio of matching query genes/all query genes to enrichment term size. We aggregated the top 100 terms (up- and downregulated) across the results for all data sets and, using rrvgo (version 1.8.0), we used Wang term semantic similarity to calculate the similarity of terms to each other using the hierarchy of GO’s graph for GO biological processes (GO:BP).(Sayols, 2020; Wang et al., 2007; Yu et al., 2010) We plotted the term enrichment similarity across the data sets with a grouping similarity threshold set to 0.80 in order to select parent terms.

### Acute Kidney Injury Signature Assessment

To assess differential expression of known acute kidney injury (AKI)-associated genes across the data sets in our study, we obtained a set of consistently differentially expressed genes associated with AKI from a previous study that aggregated 4 AKI data sets with 24-hour post-injury and time-matched controls.(Chen et al., 2020) A total of 157 and 88 genes were upregulated and downregulated, respectively, across all 4 data sets in that study providing a consensus AKI signature. Using these genes, we subset the differential expression results of the pre-cystic and cystic data sets compared to controls in our three data sets and assessed the AKI signature in each.

### Signature Reversion Searching for Drug Candidates

We mapped mouse Ensembl genes to orthologous human (GRCh38.p13) Ensembl genes and further converted them to Entrez gene IDs with BiomaRt (version 2.50.3).(Durinck et al., 2005, 2009) The disease signatures from DESeq2 were mapped to Entrez genes available in the LINCS(Subramanian et al., 2017) drug perturbation data we procured from ExperimentHub version 2.2.1 data set EH3226. Then we selected the top 100 upregulated and downregulated genes based on DEseq2’s log2 fold change as the gene signature for signature reversion (Additional files 4 and 5). This cut-off was based on the previous work of Yang et al. as described above.(Yang et al., 2022) We used the LINCS bi-directional weighted Kolmogorov-Smirnov algorithm in the signatureSearch (1.8.2)(Duan et al., 2020) package for signature querying of the LINCS drug perturbation data (EH3226) from the signaturesearchData (1.8.4)(Duan et al., 2020) package.(Subramanian et al., 2005, 2017) Briefly, this algorithm uses as input the most upregulated genes and downregulated genes as mutually exclusive lists from an experiment (in our case) as a query to then compare against the reference database of rank-transformed signatures. Similar to a weighted Kolmogorov-Smirnov statistic in gene set enrichment analysis (GSEA), the query is compared to reference lists to find if there is an overrepresentation of the reference genes at the “extreme ends” (top or bottom) of the query list.(Subramanian et al., 2005) The bi-directional method that computes a weighted connectivity score (WTCS), similar to an enrichment score (ES), calculates the representation (connectivity) by using this similarity metric to compare the degree of overrepresentation of the upregulated query genes to the top of the rank-ordered reference signature, and the overrepresentation of downregulated query genes to the bottom of the rank-ordered reference signature.(Subramanian et al., 2017) The WTCS is then scaled with values ranging from -1 (most inversely related) to 1 (most similarly related), where 0 is null (not related). Finally, the WTCS is normalized across cell lines and perturbagens within the reference database to give a normalized connectivity score (NCS).(Subramanian et al., 2017) To analyze only drugs inversely related to the ADPKD signature, we filtered results for negative NCS. Due to variability in the compound-induced expression being highly cell-line specific, we further analyzed only kidney-derived cell lines for compound results by filtering for HA1E and NKDBA compound-treated results.(Yang et al., 2022)

### Drug Candidate Prioritization

We further annotated drugs with data regarding clinical phase status, mechanism of action, and drug targets using The Drug Repurposing Hub (release 3/24/2020 https://clue.io/repurposing#download-data).(Corsello et al., 2017) The Drug Hub’s “launched” annotation refers to the highest clinical status achieved by each compound and was manually curated across multiple sources worldwide for approval (The FDA Orange Book, prescribing labels, ClinicalTrials.gov, PubMed, EU Clinical Trials Register, Canada DPD, Japan PMDA, and more).(Corsello et al., 2017) We used the “launched” status to filter potential drugs as well as the mechanism of action (MOA) and original disease indication annotations. We further prioritized by filtering for current FDA-approved drugs’ active ingredients. For this, we downloaded the Drugs@FDA database as a compressed data file (drugsatfda.zip) from the FDA website.(Center for Drug Evaluation & Research, n.d.) Compounds were filtered for active ingredients and marketing status IDs (e.g., prescription, over-the-counter, or tentative approval) regardless of form (e.g., injectible, solution, tablet, etc) (accessed August 2022).(Center for Drug Evaluation & Research, n.d.) We compared drug targets from signatureSearch to differentially expressed genes from DESeq2 results. We compiled drug target data annotations using signatureSearch from DrugBank,(Wishart et al., 2018) CLUE,(Subramanian et al., 2017) and STITCH.(Szklarczyk et al., 2016) We conducted Drug Set Enrichment Analysis (DSEA) using signatureSearch’s dsea_hyperG function with hypergeometric testing. We used the GO:MF ontology source for DSEA with gene set sizes > 10 and < 500 using the Benjamini-Hochberg procedure for multiple hypothesis correction (p-adjusted value cutoff 0.05). From the epocrates clinical database, we compiled the average monthly retail prices for drugs and black box warnings (accessed November 2022).(*Epocrates Web*, n.d.)

## Results

### Genes and Pathways Driving Cyst Progression in Mouse Models of PKD

To identify disease-associated transcriptomic signatures for ADPKD, we acquired three publicly-available *Pkd2* KO mouse data sets (Additional file 1) and performed differential gene expression analysis using DESeq2 between the ADPKD model (condition) and wild-type (control) for each data set. We identified differentially expressed genes for each data set by comparing *Pkd2* KO samples to their wild type controls, resulting in the identification of 257 differentially expressed genes for pre-cystic P70(Zhang et al., 2021), 421 for cystic P21(Lakhia et al., 2016), and 3,462 for cystic P28(Lee et al., 2019) (LFC > 1.1 or 2 and p-adjusted value < 0.05) (Figure 1A, Data files “deseq2_outputs”).(Wilk et al., 2023) The P21 and P28 cystic data sets showed excellent agreement. A total of 387 genes (92%) of the P21 differentially expressed genes were also differentially expressed in the P28 analysis. There were many more differentially expressed genes in the P28 cystic data set than the other two data sets, likely due mostly to increased detection power given there were many more P28 samples (n = 23 in cystic P28, n = 7 in cystic P21, and n = 6 in pre-cystic P70) (Additional file 1).

**Figure 1:**
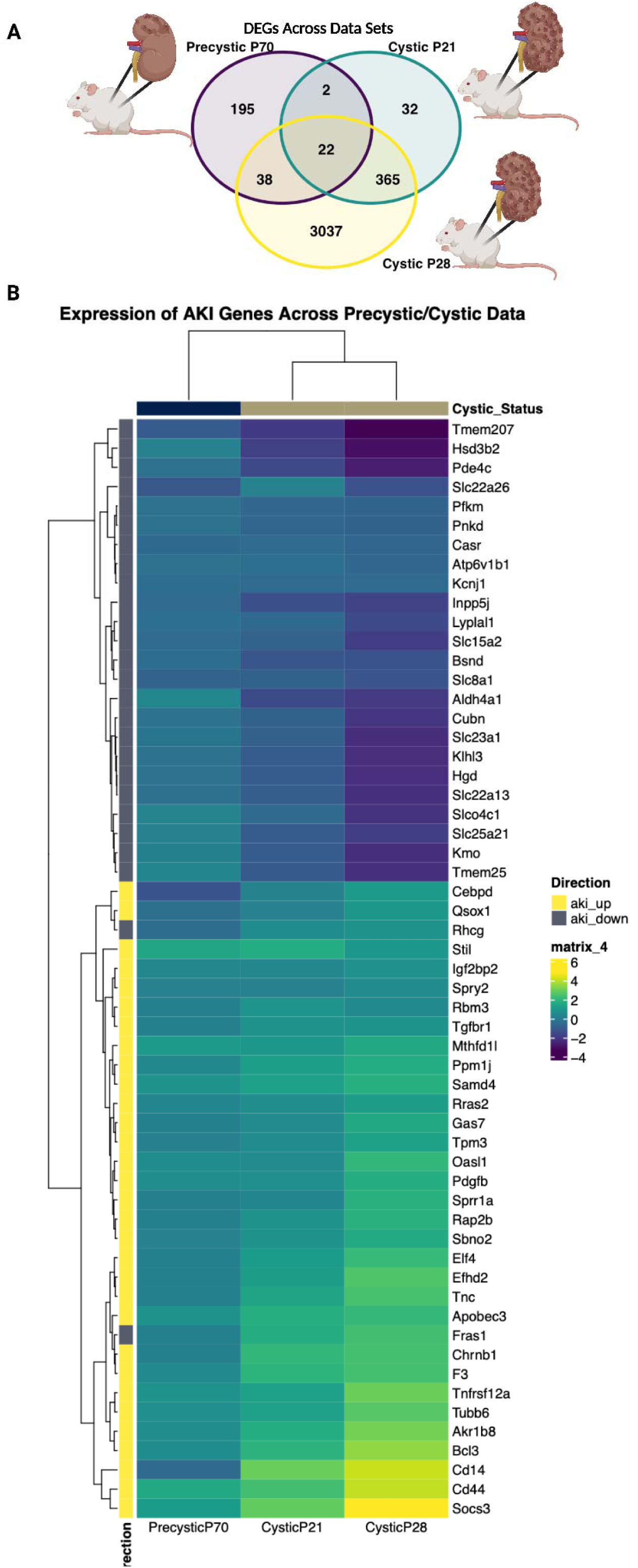
Differential Expression Analysis. A) Comparison of unique and overlapping differentially expressed genes between the 3 data sets. B) Heatmap of LFC values for AKI-associated genes for each data set.

To further assess the difference in the pre-cystic versus cystic data sets, we used the overlapping differentially expressed genes in the cystic data sets as well as the genes uniquely differentially expressed in the pre-cystic data set. We tested them for enrichment using gprofiler2 (sources Reactome, Wikipathways, GO biological process (GO:BP), GO cellular component (GO:CC), and GO molecular function (GO:MF)). There were no enrichment results for overlapping downregulated cystic and pre-cystic genes, but the upregulated genes were enriched for chemokine, immune, and inflammation pathways (Additional file 2). We noted that NAD+ pathways were also enriched for the overlapped upregulated cystic genes. Due to NAD+ regeneration increase in response to high glycolysis, we hypothesize that this may be evidence of the Warburg-like effect previously seen in ADPKD.(Podrini et al., 2020; Rowe & Boletta, 2014) Only those genes that were overexpressed compared to wild type in pre-cystic, but not in either cystic data sets, were largely involved in cell division and cell cycle pathways (Additional file 3). Furthermore, enrichment of differentially expressed genes unique to the pre-cystic data set were also *Cdk1* pathway members which, in particular, have been indicated as a critical driver of cyst proliferation in ADPKD (Additional file 2).(Zhang et al., 2021)

Due to the cystic-associated differential expression being enriched for excessive immune, inflammatory, and keratin upregulation and prior literature supporting that acute kidney injury (AKI) can accelerate PKD in mouse models, we sought to compare AKI-associated genes across all 3 data sets.(Kurbegovic & Trudel, 2016) For this, we obtained a list of consistently upregulated and downregulated genes from previous work by Chen et al., where 4 AKI data sets with 24 hour post-injury and time-matched controls were analyzed for consistent differential expression.(Chen et al., 2020) They identified a total of 157 upregulated genes and 88 downregulated genes across all 4 data sets. We then used these genes to subset the differential expression results of the pre-cystic and cystic data sets compared to controls (> 0.15 LFC and < 0.1 p-adjusted value). The P21 and P28 cystic gene expression clustered together and the known upregulated AKI genes had upregulated LFCs in the cystic data sets, and downregulated AKI genes had downregulated LFCs in the cystic data sets (Figure 1B). We conclude that the AKI molecular phenotype was much stronger in the cystic data compared to the pre-cystic data suggesting that it may confound the PKD molecular phenotype.

### Inflammatory Pathways Driving Variation in ADPKD Cystic Transcriptomic Signature

We then asked what gene pathways and modules were overrepresented in pre-cystic and cystic disease signatures, and how they differ from the full differentially expressed gene list. These disease signatures consisted of the 100-200 differentially expressed genes from each data set which mapped to LINCS measured or inferred genes as Entrez gene IDs (see Methods and Additional files 4 and 5). We found our signatures to be consistent with previous ADPKD functional enrichment analyses, including chemokine activity/receptor binding, and dysregulation of multiple metabolic pathways (Figure 2A-C, Data files “fea”) (p-adjusted value < 0.05, sources GO, Reactome, and WikiPathways).(D. Liu et al., 2019; Malas et al., 2020; Wilk et al., 2023) Consistent with the literature, both of the cystic signatures (P21 and P28) were enriched for multiple chemokine and interleukin pathways (Figure 2B-C), and the pre-cystic signature was enriched for multiple cell cycle and cell division pathways (Figure 2A). Pre-cystic transcriptomic signature resulted in NIMA Kinases (NEKs) as a top enrichment term. NEKs are characteristically dysregulated in PKD, which are involved in cilia assembly and function and may further play a role in PKD via cell cycle and mitotic spindle dysregulation.(Fry et al., 2012) Regulation of arterial blood pressure and blood circulation was downregulated in the pre-cystic signature, a previously noted indication of hypertension (Figure 2A).(Chapman et al., 2010) Hypertension is a common comorbidity of ADPKD, likely due to the many complex overlapping roles between polycystin 1 and 2 and vascular structure and function. Impaired polycystin 1 and 2 function leads to decreased nitric oxide release and synthesis, which then leads to renin-angiotensin-aldosterone (RAAS) activation and cyst growth.(Chapman et al., 2010) Furthermore, cyst growth further perpetuates hypertension via RAAS, accounting for some of the strong comorbidity between ADPKD and hypertension.(Chapman et al., 2010) Hypertension is a driving factor in ADPKD for end-stage renal disease (ESRD), so it is intriguing that blood pressure regulation pathways were only identified in the pre-cystic signature and not the cystic signatures.

**Figure 2:**
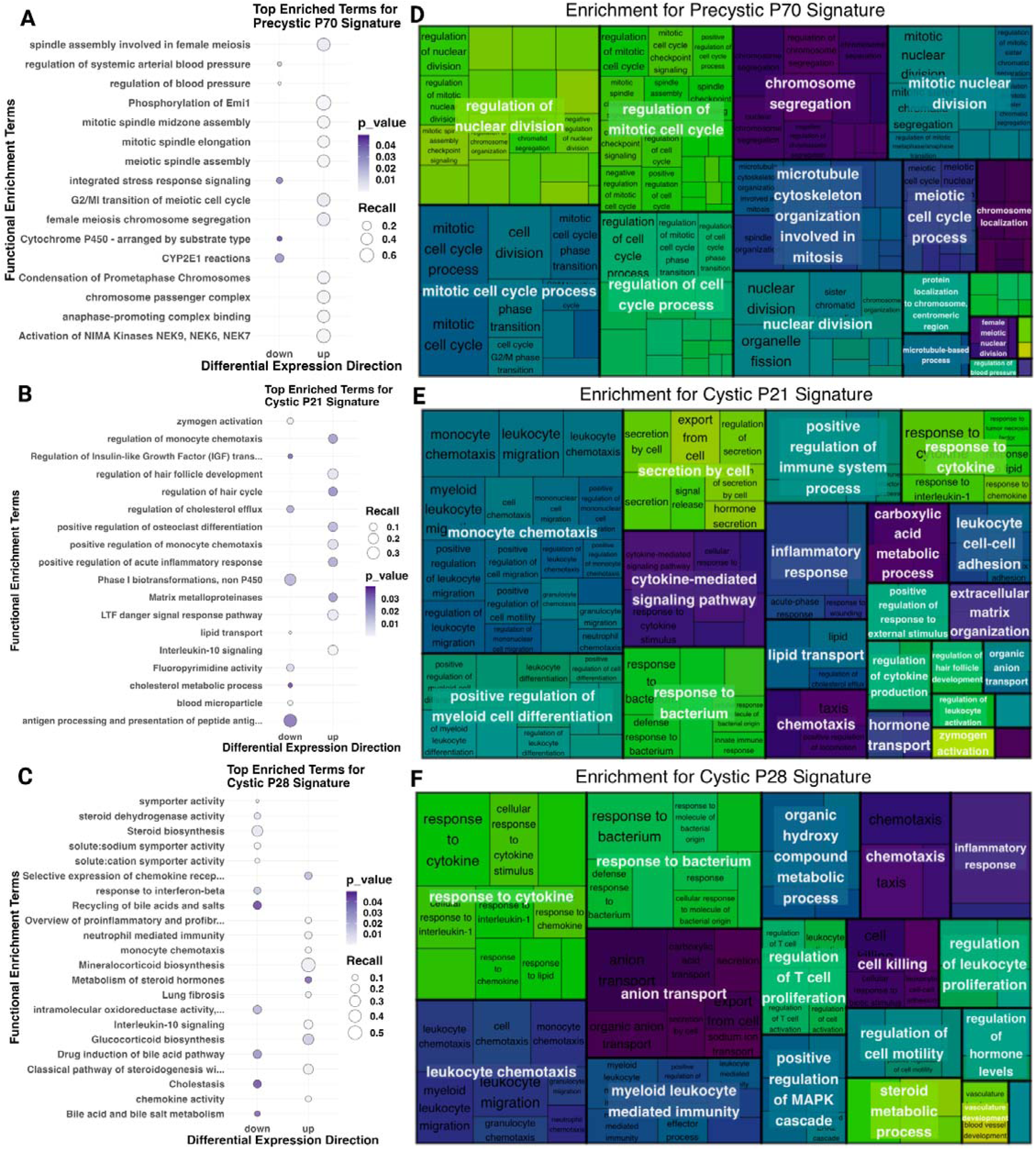
Pathway Enrichment of Pre-cystic and Cystic Transcriptomic Signatures. Bubbleplots of pathway enrichment using GO, Reactome, and WikiPathways of upregulated and downregulated genes from the A) pre-cystic P70 transcriptomic signature, B) cystic P21 transcriptomic signature, and C) cystic P28 transcriptomic signature, with point size dependent on the number of query genes matching to each term and color dependent on the p-value. Treemaps display the enrichment hierarchy, using Wang similarity to reduce GO:BP term redundancy with parent terms consisting of largest boxes and enriched child terms within for each signature, and varying child terms reflecting term size for D) pre-cystic P70 signature E) cystic P21 transcriptomic signature, and F) cystic P28 transcriptomic signature.

To further resolve terms and discover parent pathway terms, we sought to reduce GO:BP enrichment results by considering the GO term hierarchical system using Wang similarity. The -log10 of the p-values were used as scoring for terms with a similarity threshold of 0.8, allowing us to visualize the nested enrichment hierarchy (Figure 2D-F). We, therefore, identified the most representative parent terms for the pre-cystic signature as cell cycle and cell division (Figure 2D), and the most representative parent terms for the cystic signatures as inflammatory response, cytokine response, and leukocyte activity (Figure 2E-F). One particularly interesting parent term that was consistent between the cystic signatures was vascular development. As previously mentioned, as renal cysts progress in ADPKD, vascularization of cysts increases and further exacerbates inflammation and other immune responses, somewhat mimicking AKI.(Zimmerman et al., 2020) This, as well as our comparison of AKI genes in cystic differentially expressed genes (Figure 1B), supports the hypothesis that the already cystic disease signatures may be confounded by excessive inflammation and disease response, potentially masking some critical driving mechanisms of ADPKD pathogenesis.

### Signature Reversion of the Pre-Cystic PKD Profile

We decided to focus on the pre-cystic P70 transcriptomic signature for signature reversion drug repurposing candidate prioritization, based on our findings that it may provide the strongest biological signals for ADPKD driving mechanisms, compared to the cystic signatures which may be confounded by inflammation and PKD pathogenic response (Figures 1-2). To identify drugs that might reverse the cellular PKD phenotype, we applied signature reversion analysis, filtering for drug signatures with negative normalized connectivity scores (NCS) (drug signatures inversely related to the pre-cystic signature) in kidney-derived cell lines. This analysis resulted in 730 drug candidates. We then filtered for “launched” drugs via the Drug Repurposing Hub, narrowing down to 178 drug candidates. Finally, we filtered for FDA-approval status of drug candidates by comparing the active ingredients with either over-the-counter, prescription, or tentative-approval statuses as of August 2022. This resulted in 109 candidates (Data file “sigsearch_outputs”).(Wilk et al., 2023) The most frequent mechanisms of action (MOAs) were in line with previously studied drug MOAs for ADPKD (i.e., topoisomerase inhibitors, tubulin interfering agents, and dopamine receptor antagonists) (Figure 3A).(Asawa et al., 2020; Paul et al., 2019; Woo et al., 1994) For the drugs with original disease area indication available, we compared the most frequent original indications for our 109 FDA-approved drug candidates (Figure 3B). While it was unsurprising to observe oncological/malignancy drugs as a frequent repurposing option for ADPKD due to the aforementioned overlap of ADPKD and cancer, the unexpected most frequent original disease indication was neurology/psychology.(Seeger-Nukpezah et al., 2015)

**Figure 3:**
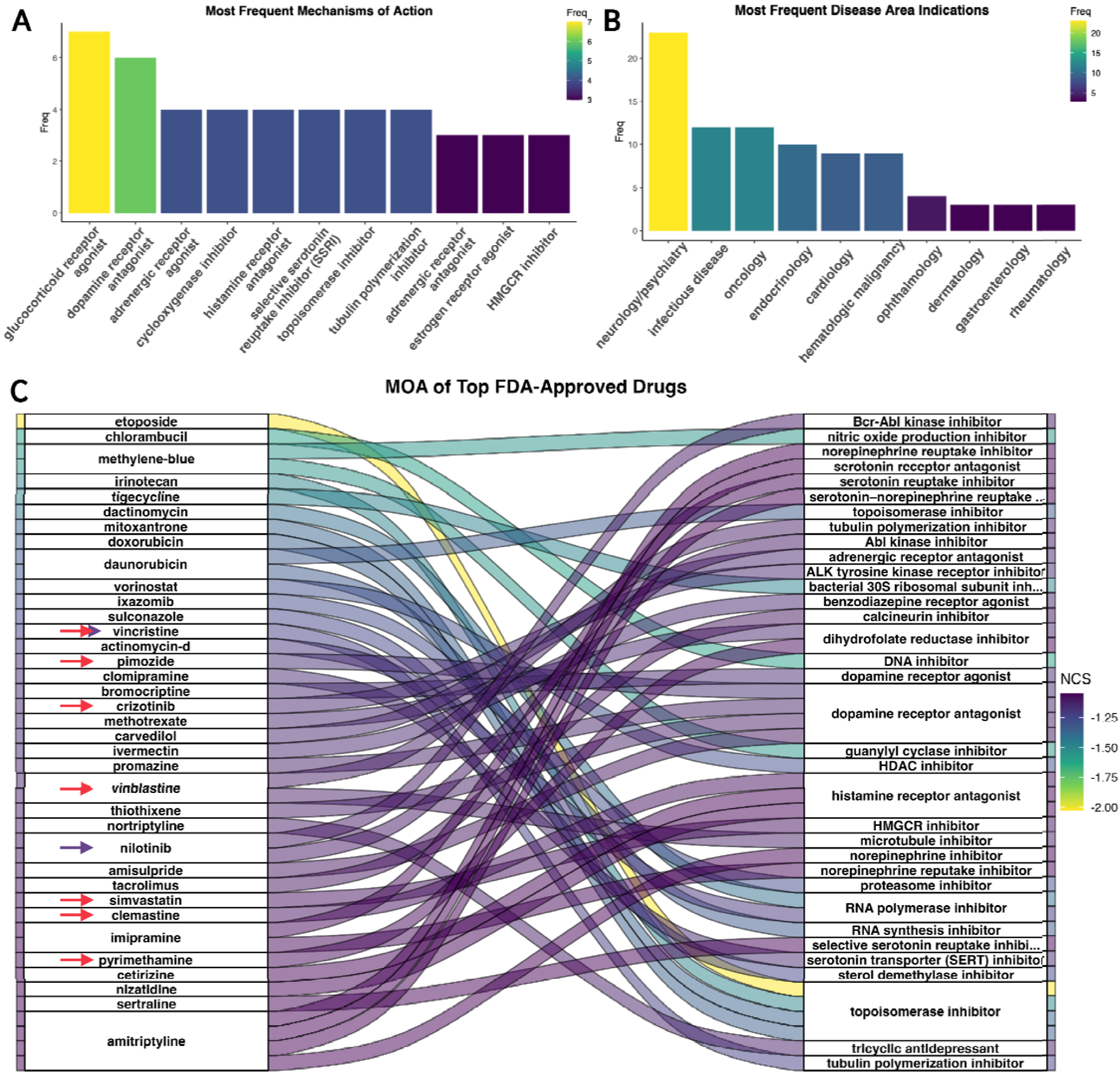
FDA-Approved Drug Candidates MOA and Original Indications from Pre-Cystic Signature Reversion. A) The most frequent MOA of all FDA-approved candidates, B) the most frequent disease areas for original indications of candidates, and C) alluvial plot of the top FDA approved candidates by NCS and their MOA, with drugs that have been previously explored for PKD treatment marked with red arrows, and drugs previously explored for CKD and/or AKI are marked with dark purple arrows.

When further investigating the top individual drugs by sorting for the most inversely enriched NCS, we found multiple drugs previously investigated as repurposing candidates for ADPKD (red arrows in Figure 3C) and/or AKI or chronic kidney disease (CKD) (purple arrows in Figure 3C). For plotting purposes, we selected the top 35. Our top candidates by NCS that were previously studied in the literature for PKD included vincristine, clemastine, simvastatin, pyrimethamine, pimozide, crizotinib, and vinblastine, and previously studied treatments for chronic kidney diseases and AKI included vincristine and nilotinib.(Fontecha-Barriuso et al., 2018; Gilbert et al., 2011; Iyoda et al., 2011; Strubl et al., 2020; Tajti et al., 2020; Takakura et al., 2011; van Dijk et al., 2001; Woo et al., 1994)

### Pre-cystic Signature Drug Candidate Target Expression in Cystic Kidney Tissue

We then asked if the drugs we uncovered by signature reversion of the pre-cystic data set have known drug targets with upregulated gene expression in the pre-cystic as well as the cystic data sets. These drugs would therefore be ideal candidates for patients with already cystic kidneys as well. Drug targets for the 109 FDA-approved drug candidates from pre-cystic signature reversion were compared against upregulated differentially expressed genes for each data set, revealing even more targets upregulated in the cystic data sets P21 and P28 (Figure 4A-B) than the pre-cystic data set from which the candidates were found (Figure 4A). Total upregulated genes matching to unique drug candidates were 12, 16, and 38 for data sets P70, P21, and P28, respectively. We further investigated the drugs with robust cystic target expression by prioritizing drugs with upregulated drug targets in both cystic data sets, leaving us with a total of 16 drugs targeting 29 unique cystic-upregulated genes (Figure 4C, Table 1).

**Figure 4:**
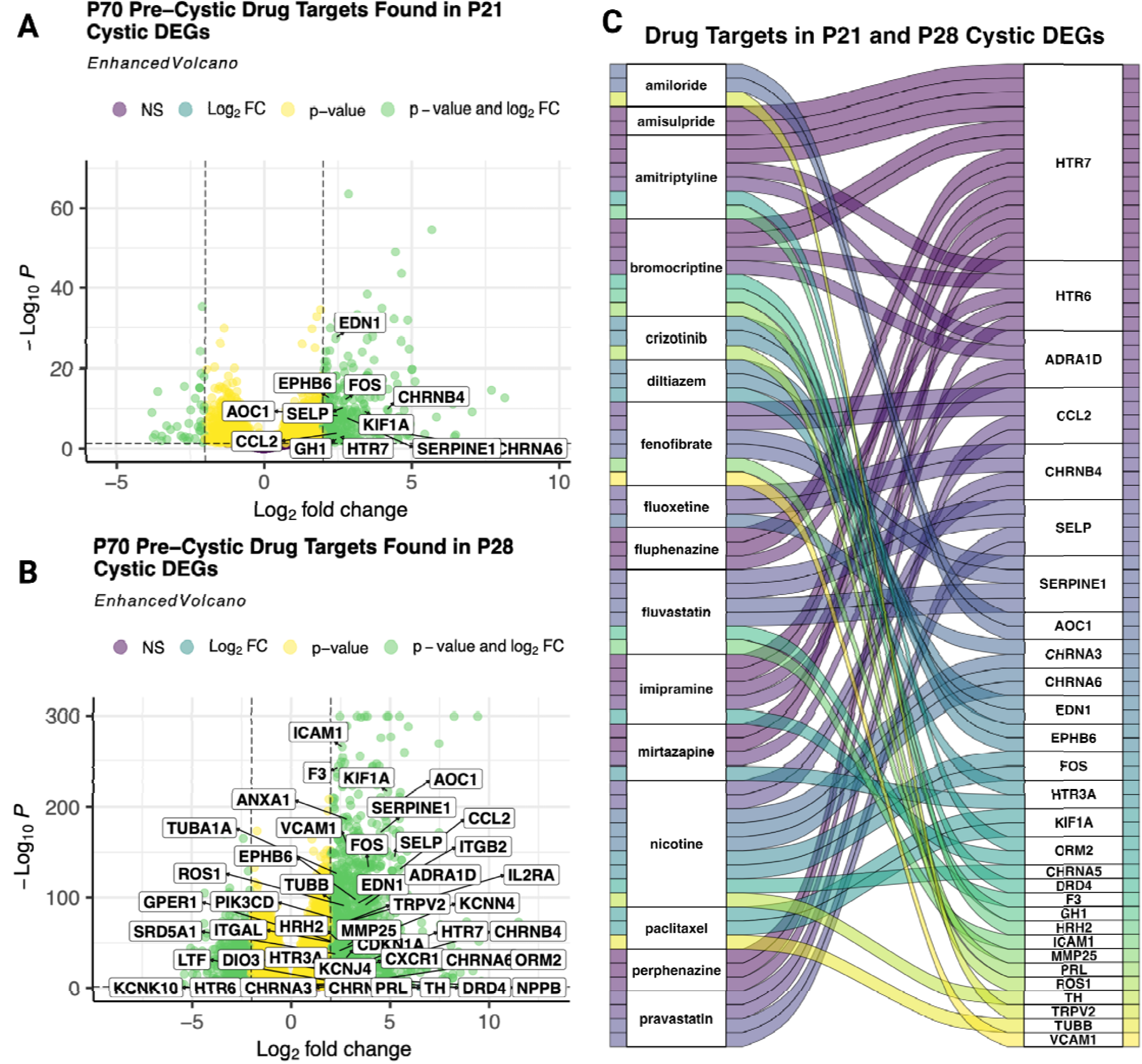
Pre-cystic Signature Reversion Drug Candidates Target Upregulated Genes in Pre-cystic and Cystic Profiles. Volcano plots of differentially expressed genes (p-adjusted < 0.05) with differential expression of candidate drug targets labeled in the A) P21 cystic (LFC > 2.0), and B) P28 cystic (LFC > 2.0). C) Alluvial plot of the 16 drugs with upregulated drug targets in both cystic data sets connected to their targets that were also found upregulated in the cystic differentially expressed genes (A and B).

**Table 1:**
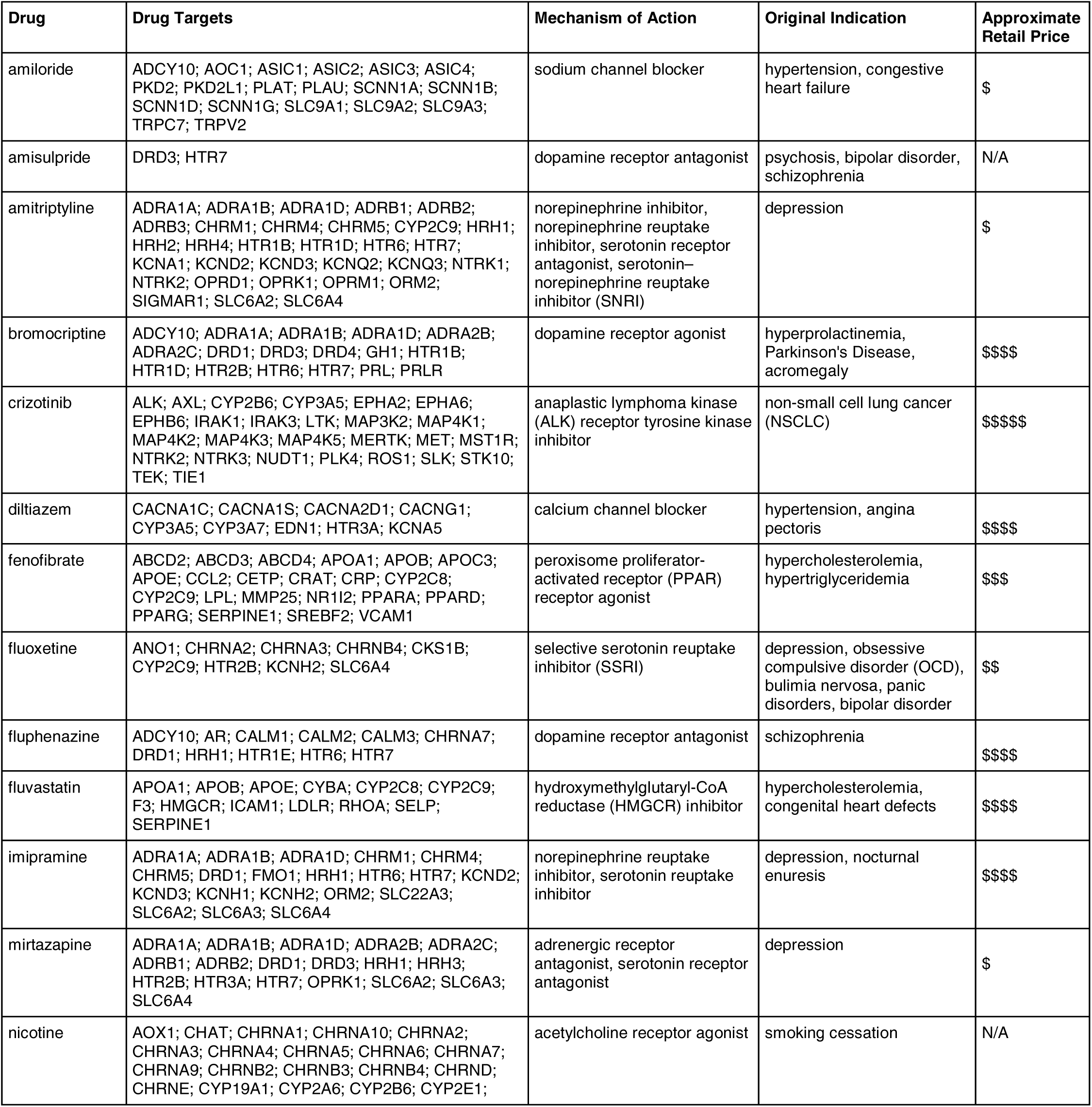
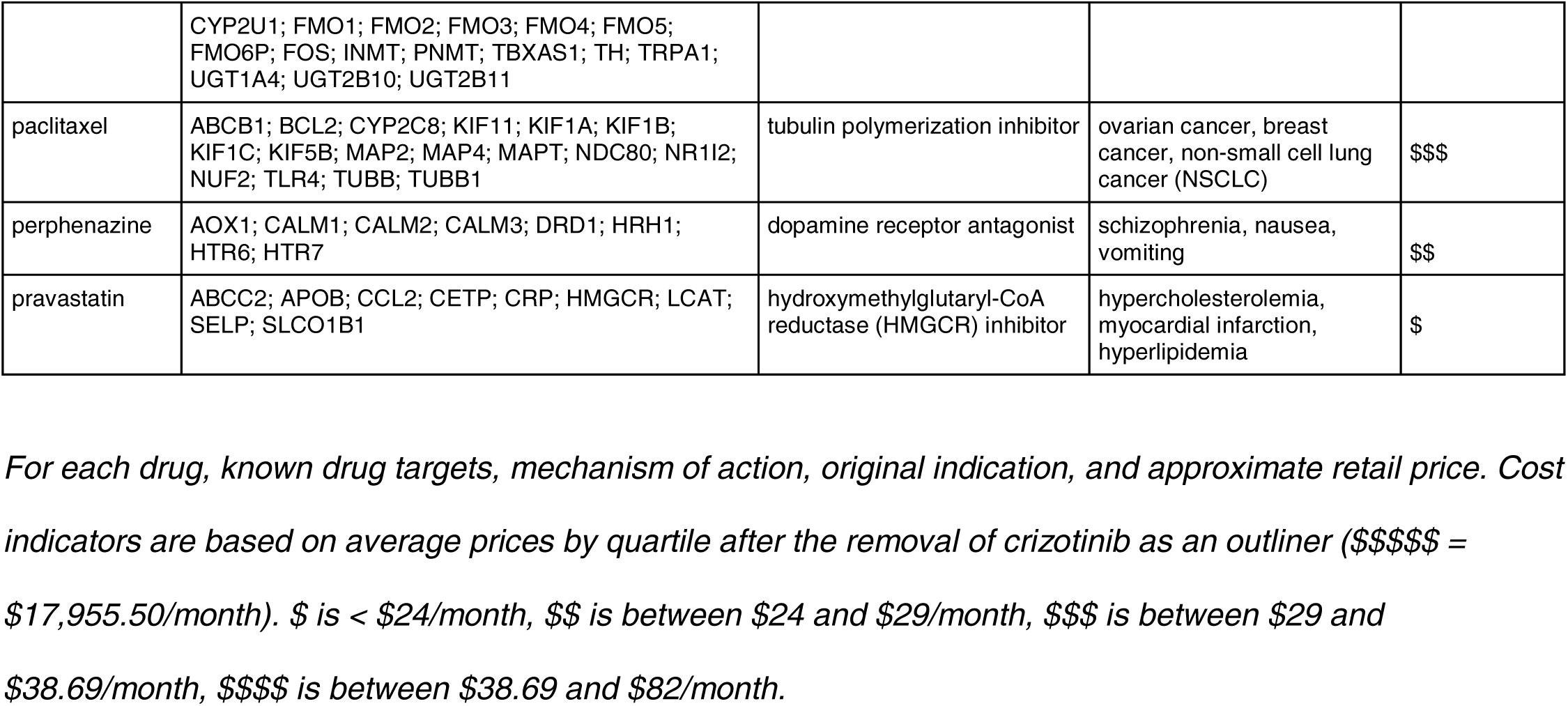
Summary of Prioritized Drug Candidates.

Table 1: Summary of Prioritized Drug Candidates.

Located after the ‘Conclusions’ section.

### Pathway and Drug Set Enrichment Analysis of Top Drug Repurposing Candidates

We further considered our 16 prioritized drug candidates by investigating pathways for which they were enriched and the drug-target networks within those enriched pathways. For this, we used signatureSearch’s drug set enrichment analysis (DSEA) with hypergeometric testing (p-adjusted value < 0.05) and plotted the top enriched GO:MF terms by p-adjusted value (Figure 5A). Several of the top resulting pathways are known to be involved in PKD, including oxidoreductase activity, G protein-coupled serotonin receptors, and histamine receptor activity.(Holditch et al., 2019; Sudarikova et al., 2021; Trudel et al., 2016) Closer inspection of the protein interactions of our prioritized drugs for oxidoreductase activity showed drug metabolism activity specifically with the genes encoding cytochrome P450 enzymes, *CYP2C8* and *CYP2C9* (Figure 5B). *CYP2C8* and *CYP2C9* P450 enzymes also metabolize fatty acids, a metabolic pathway well known to be altered in ADPKD disease progression.(Podrini et al., 2020) *CYP2C8* and *CYP2C9* metabolize many other drugs and are highly polymorphic, heavily impacting pharmacokinetics.(Neuvonen et al., 2008) As shown here, *CYP2C8* metabolizes paclitaxel, fluvastatin, and fenofibrate, and *CYP2C9* also metabolizes fluvastatin and fenofibrate as well as amitriptyline and fluoxetine (Figure 5B). It should be noted that these CYP interactions are not the only CYP interactions for our prioritized drugs, but are part of the drug-target network for this GO:MF oxidoreductase activity pathway. For example, fluoxetine also interacts with *CYP2C19* and *CYP3A4* but is most metabolized by *CYP2D6*, to which it is also a strong inhibitor.(Deodhar et al., 2021)

**Figure 5:**
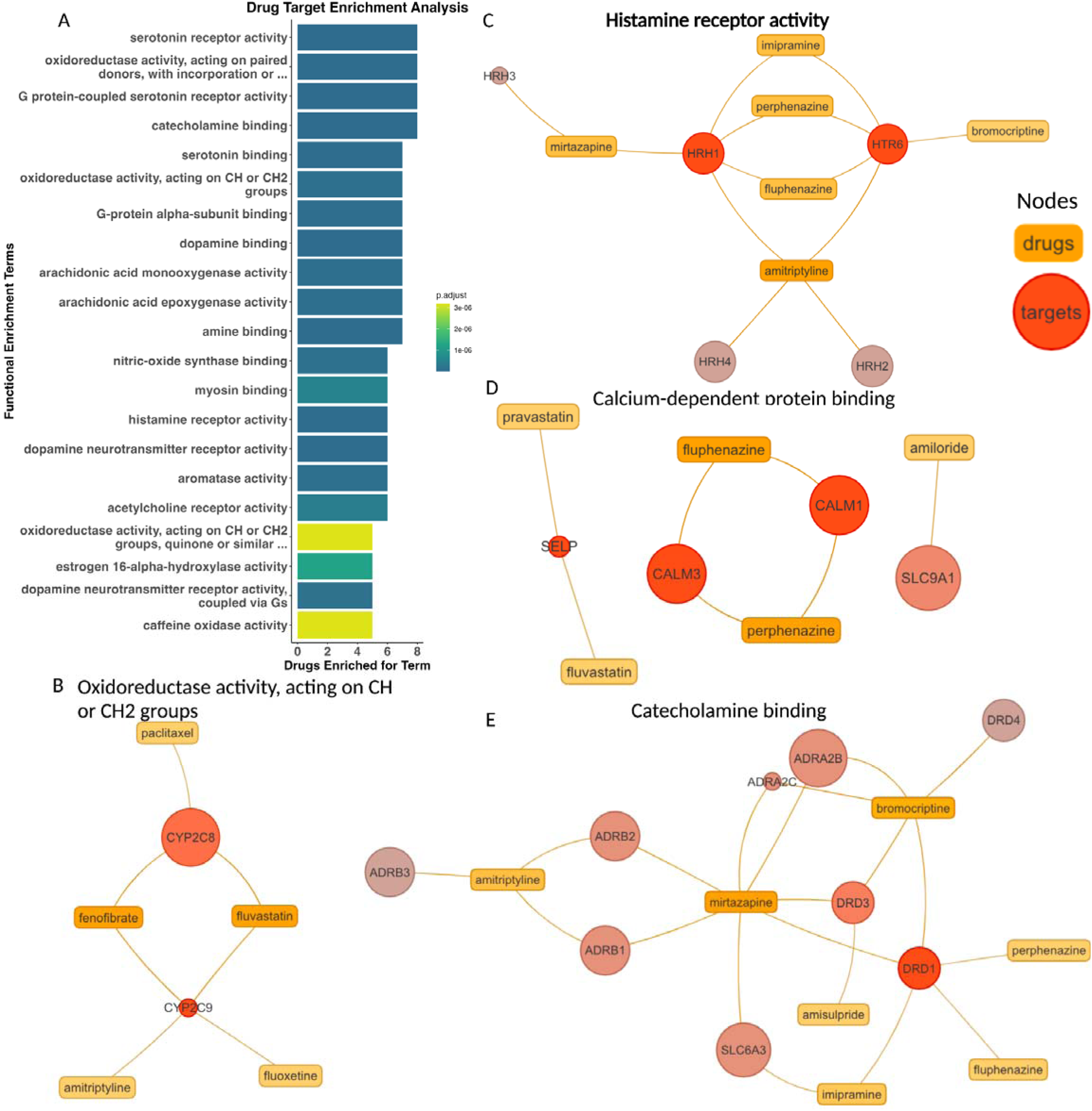
Prioritized Drug Targets’ Pathways and Binding. Top enriched GO:MF pathways of targets from 16 prioritized drugs by p-adjusted value as A) a barplot and B-E) specific pathways shown as networks of prioritized drugs (orange rectangular nodes) and their targets (red circular nodes) with color shade defined by the number of connections to other nodes.

Drug binding with histamine receptors showed mirtazapine, a tetracyclic antidepressant, also targets *HRH3* and *HRH1* (Figure 5C). Mirtazapine has strong antihistamine effects and was found to be well-tolerated in CKD patients for the treatment of chronic kidney disease-associated pruritus (CKD-aP).(Mehrpooya et al., 2020) Another drug originally indicated for depression that was also shown to have antihistamine effects is amitriptyline. Amitriptyline has already been investigated for repurposing in osteoarthritis and gout for its anti-inflammatory properties.(Franco-Trepat et al., 2022; Wluka et al., 2021)

Fluphenazine and perphenazine, two other antidepressants, were both additionally shown to interact with histamine receptors and, further, bind to calmodulin (encoded by *CALM1* and *CALM3*) as calcium-dependent protein binding (Figure 5C-D). Calmodulin is a primary calcium sensor, likely playing a critical role in PKD pathogenesis in response to decreased calcium levels due to polycystin 1 or polycystin 2 defect, leading to increased levels of cAMP.(Chebib et al., 2015) Furthermore, fluphenazine’s calmodulin inhibition has been hypothesized to prevent fibrosis via calcium-saturated calmodulin-dependent myosin light chain kinase (CaM-MLCK) as well as cancer cell invasion and metastasis.(Levinson et al., 2004; Villalobo & Berchtold, 2020)

The catecholamine binding pathway enriched in our prioritized drug targets includes binding to dopamine receptors Drd1 and Drd3 (Figure 5E). As we previously discussed, dopamine receptor antagonists are promising for ADPKD treatment, and have been shown to reduce cystic growth and cell proliferation in *Pkd1*-/- mice.(Paul et al., 2019) HDAC5 export maintains renal epithelial architecture via de-repression of MEF2C target genes and is dependent on polycystin calcium influx. Dopamine receptor antagonists, specifically for Drd3, have been shown to restore HDAC5 nuclear export in *PKD1loxP*/*loxP* cells.(Paul et al., 2019) Bromocriptine, amisulpride, and mirtazapine were all shown to target Drd3 specifically, and therefore may be particularly effective in cyst reduction and cell proliferation by HDAC5 nuclear export. This shows that our prioritized candidates target pathways with known involvement in ADPKD pathogenesis.

## Discussion

We analyzed publicly available *Pkd2* KO mouse RNA-Seq data sets to assess differential gene expression, pathway enrichment, and transcriptomic signature reversion to identify potential drug repurposing candidates to identify prioritized therapeutics for the ADPKD community. Our work extends beyond the original analyses reported for these data sets and highlights the important role public data repositories can have in developing therapeutic paths for ADPKD. Additionally, as the P70 pre-cystic data set consisted of tissue collection at the pre-cystic/cyst-initiation stage, this presented an opportunity to identify early molecular changes associated with cystic kidney disease and potentially confounding signatures associated with full cystogenesis (i.e., fibrosis and inflammation).(Zhang et al., 2021) This is ideal for prioritizing drug repurposing candidates that may be effective for early or preventative treatment. Further, by comparing the drug targets for the pre-cystic P70 candidates to differentially expressed genes of the P21 and P28 PKD mouse models (*Pkhd1*-cre;*Pkd2*^F/F^), we identified 29 unique drug targets that were differentially expressed in both *Pkd2* cystic data sets for 16 drug candidates. These indicate drug candidates that may effectively treat pre-cystic as well as cystic ADPKD.

Consistent with previous findings, we identified genes known to have a significant role in the pathogenesis of ADPKD including *WNT7A, LCN2,* and *MMP7* (Additional files 4 and 5; Data files “deseq2_outputs”).(A. Li et al., 2018; Petra et al., 2022; Viau et al., 2010; Wilk et al., 2023; D. Zhou et al., 2017) The previously noted biomarkers of CKD progression *LCN2* and *MMP7* were upregulated in both P28 and P21 cystic data sets.(Petra et al., 2022; Viau et al., 2010; D. Zhou et al., 2017) In line with other studies, most of the upregulated pathways in the cystic data sets P21 and P28 were associated with inflammation, leukocyte activity, NAD+ activity, and cytokine response. Additionally, genes differentially expressed only in the P70 pre-cystic data set were enriched for other pathways characteristic of ADPKD pathogenesis, including upregulation of cell cycle and *Cdk1*, a known driver of cyst proliferation in ADPKD.(Zhang et al., 2021) We further found that the P21 and P28 cystic differentially expressed genes were highly similar to consistently upregulated and downregulated genes in AKI compared to the P70 pre-cystic differentially expressed genes (Figure 1B). Finally, while gene expression changes do not always reflect disease etiology and may only be due to downstream disease responses, careful selection of both the data sets (such as the pre-cystic P70) and drug candidates is critical.(Porcu et al., 2021) For example, some of our top candidates, such as phosphodiesterase inhibitors (Figure 3A), are predicted to further increase cellular cAMP, which may actually exacerbate ADPKD-associated cellular phenotypes due to decreased intracellular mechanisms from defective PC1/PC2 calcium-sensing and transport. This further underscores the need for placing such analyses in the biological context.

In this study, we prioritized drug repurposing candidates by selecting FDA-approved compounds (i.e., those that already have safety profiles for other diseases) predicted to reverse ADPKD transcriptomic signatures so they could be most greatly expedited as potential ADPKD treatments. While previous drug screens have identified ADPKD drug repurposing candidates using *pkd2* mutant tail phenotypes in zebrafish (Cao et al., 2009; Metzner et al., 2020) and in *Pkd1* knock-out mice across disease stages (Malas et al., 2020), our study is the first to our knowledge to apply signature reversion to the problem of identifying drug repurposing candidates for ADPKD.(J. X. Zhou & Torres, 2023) While our approach, identified the top candidates from these previous studies with our original pre-cystic signature reversion results, they were subsequently filtered out during our prioritization schema because they are not identified as launched, FDA approved, or did not have overexpressed drug targets in the cystic profiles. By further investigating each drug’s known mechanism of action and their expressed or differentially expressed drug targets in all 3 data sets (i.e., P70, P21, and P28) we identified additional drugs previously investigated for ADPKD treatment in clinical trials (pravastatin) as well as other preclinical studies (e.g. fenofibrate, paclitaxel), and uncovered promising novel candidates. For example, topoisomerase inhibitors and tubulin interference drugs were previously identified for their ability to reduce cyst growth in a large screen of ∼8,000 compounds in *Pkd1*-null 3D-grown cells.(Asawa et al., 2020) Topoisomerase inhibitors, originally indicated for cancer, disrupt the ability of cells to repair single-stranded DNA breaks incurred during DNA replication.(“Topoisomerase Inhibitors,” 2020) As topoisomerase activity is drastically increased in cancer due to rapid division (as is likely the case in ADPKD given the overlapping molecular characteristics between cancer and ADPKD(Seeger-Nukpezah et al., 2015)), inhibiting the repair of single-stranded nicks from DNA replication leads to replication inhibition followed by cell death.(“Topoisomerase Inhibitors,” 2020) Notably, tubulin interference drugs were previously found not only to reduce cyst growth in 3D-grown cells, but also to reduce growth without impacting cell viability.(Asawa et al., 2020) Interference of tubulin dynamics has been previously studied in ADPKD mouse models, resulting in significantly increased survival time, renal function, adult body weight, and decreased cyst growth.(Woo et al., 1994) However, another class of drugs, dopamine receptor antagonists, are under-researched in ADPKD even though they increase *HDAC5* nuclear export.(Paul et al., 2019) In healthy kidney cells, the polycystin complex formed by the *PKD1* and *PKD2* proteins facilitates the nuclear export of *HDAC5*, resulting in the maintenance of renal epithelial architecture.(Paul et al., 2019) Furthermore, *Pkd1*-/- mice treated with dopamine receptor antagonists did exhibit cyst reduction and increased body weight and activity.(Paul et al., 2019)

However, the most frequent MOA of these FDA-approved drug candidates was glucocorticoid receptor agonists (Figure 3A). While, to our knowledge, these have not been previously investigated for ADPKD, there are multiple, intriguing potential connections between glucocorticoid receptor agonists and ADPKD. Glucocorticoids are steroid hormones that are often used for inflammation and immune suppression across multiple diseases by repressing the activity of immunologic transcription factors such as nuclear factor kappa-light-chain-enhancer of activated B cells (NF-κB), AP-1, and T-bet.(Hudson et al., 2018) Additionally, glucocorticoids have more recently been investigated for potential pathway involvement in nephrological diseases and in pathways such as Wnt signaling and autophagy.(Przybyciński et al., 2021) While neurology/psychology-indicated drugs do have the most compounds annotated in the Drug Repurposing Hub we used for annotation further investigation is warranted, especially due to the noted comorbidity between ADPKD and depression.(Simms et al., 2016)

Additionally, the HDAC inhibitor vorinostat has been found to treat mechanisms similar to ADPKD pathogenic mechanisms, including increased renal size, extracellular matrix accumulation, and increased cell proliferation.(Seeger-Nukpezah et al., 2015) Vorinostat treatment in diabetic neuropathy rat models have resulted in decreased renal size while similar treatment in a renal epithelial cell (NRK52-E) resulted in decreased extracellular matrix expression and cell proliferation.(Gilbert et al., 2011; N. Liu & Zhuang, 2015; Mishra et al., 2003) In general, HDAC inhibitors have been of interest in treating ADPKD. Multiple studies in ADPKD pre-clinical models showed reduced cyst formation, cyst growth, improved renal function, and decreased cell proliferation following HDAC inhibition.(N. Liu & Zhuang, 2015) Interestingly, our initial top FDA-approved drugs (by most negative NCS) included three drugs that have been found to inhibit STAT family genes (Figure 3C). One study investigating STAT-targeting drugs for ADPKD therapies suggested pimozide inhibits STAT5, crizotinib inhibits STAT3 phosphorylation, and pyrimethamine inhibits STAT3 by dimerization.(Strubl et al., 2020) Pyrimethamine, an antiparasitic compound, has been further tested in ADPKD preclinical models and was found to decrease cell proliferation in ADPKD patient-derived cultured epithelial cells and prevent renal cyst formation in *Pkd1*-KO mice.(Takakura et al., 2011)

Another FDA-approved candidate, the tyrosine kinase inhibitor nilotinib, was previously named as a top candidate for the treatment of CKD in a study that combined gene expression data across 8 different glomerular diseases that lead to CKD using a different signature reversion method.(Tajti et al., 2020) Additionally, when studied in rats for CKD, nilotinib-treated rats had preserved renal function, decreased profibrotic gene expression, and significantly prolonged survival.(Iyoda et al., 2011)

We further compared the targets of our FDA-approved drug candidates from pre-cystic P70 signature reversion to differentially expressed genes from the cystic data sets P21 and P28. Among the 16 FDA-approved candidates with known drug targets overexpressed in both cystic data sets, there were 3 drugs currently in clinical trials, or in the same class of drugs in clinical trials, for ADPKD. For example, pravastatin is already in phase 4 clinical trials for ADPKD and phase 2 clinical trials in combination with acidosis treatment for ADPKD patients with worsening CKD (ClinicalTrials.gov Identifiers: NCT03273413 and NCT04284657).(Cadnapaphornchai et al., 2014) Another prioritized candidate, fluvastatin, belongs to the same drug class as pravastatin: cholesterol-lowering drugs 3-hydroxy-3-methyl-glutaryl coenzyme A (HMG-CoA) reductase inhibitors, referred to as statins. Statins have been studied for the treatment of ADPKD since 1995 due to their anti-proliferative, anti-inflammatory, and antioxidant properties.(Ecder, 2016; Gile et al., 1995; Namli et al., 2007) Finally, amiloride is being tested in clinical trials for the prevention of hypertension in ADPKD patients as hypertension is a common comorbidity of ADPKD (ClinicalTrials.gov Identifier: NCT05228574).(Chapman et al., 2010; Seeger-Nukpezah et al., 2015)

Other prioritized candidates included previously studied ADPKD therapeutics in preclinical models. Fenofibrate is an agonist of peroxisome proliferator-activated receptor α (PPARα), which promotes fatty acid β-oxidation and oxidative phosphorylation, both pathways decreased in ADPKD.(Lakhia et al., 2018) Further, treatment of *Pkd1RC/RC* mice with fenofibrate resulted in reduced renal cyst volume, proliferation, and infiltration of inflammatory cells as well as reduced bile duct cysts and liver inflammation and fibrosis.(Lakhia et al., 2018) Additionally, paclitaxel has been studied as a potential repurposing candidate for ADPKD given its ability to arrest the cell cycle, and we identified multiple targets of paclitaxel upregulated in both cystic data sets (*KIF1A* and *TUBB*, Figure 4D).(Nguyen et al., 2021)

Due to ADPKD being a chronic condition, it is also important to consider long-term adverse drug events as well as cost when further prioritizing candidates. For retail cost associations, we manually compared each drug price from the epocrates clinical database and found the average monthly cost was less than $83 (after removing one outlier, crizotinib, with an average monthly cost of $17,956) and listed the average monthly cost by quartile (Table 1). We can further de-prioritize other drugs due to their long-term adverse events and side effects specifically unfit for patients with kidney disease, including paclitaxel, amiloride, and nicotine. Amiloride has a black box warning for increased risk of hyperkalemia in patients with renal impairment.(*Amiloride/hydrochlorothiazide Black Box Warnings*, n.d.) Nicotine is particularly unfit for kidney disease patients as it has been linked with increased renal injury severity in kidney disease preclinical models and potentially leads to increased reactive oxygen species in pro-fibrotic pathways.(Jain & Jaimes, 2013) Lastly, paclitaxel has multiple severe adverse events as black box warnings, and as such may not be practical long-term treatment options.(*Paclitaxel Black Box Warnings*, n.d.)

Based on their predicted ability to reverse ADPKD molecular phenotypes, safety profiles, cost, and novel implication in ADPKD, we suggest bromocriptine and mirtazapine as prioritized ADPKD drug repurposing candidates. Bromocriptine had one of the top most negative NCSs, suggesting the greatest anti-relation to pre-cystic gene expression (Figure 3C), had multiple targets upregulated in cystic data sets (P21 and P28) (Figure 4D), and also targets Drd3, which was previously found as a potential target for restoring HDAC5 nuclear transport in ADPKD (Figure 5E).(Paul et al., 2019) Bromocriptine directly impacts other ADPKD pathogenic pathways, specifically by decreasing cAMP by adenylyl cyclase, as well as MAPK inhibition by blocking phosphorylation of p42/p44 MAPK.(Lu et al., 2019) Furthermore, bromocriptine had fewer and less severe reported adverse drug events and, to our knowledge, no black box warnings.(*Bromocriptine Adverse Reactions*, n.d.) We further highlight mirtazapine, which has an average monthly cost of less than $24 and, like bromocriptine, also has antihistamine effects and targets Drd3 (Figure 5E). Mirtazapine, like most other antidepressants, has a black box warning for the risk of suicide, especially in children and young adults, but has generally been found to be well tolerated.(Carvalho et al., 2016; Friedman, 2014)

Limitations of this current study include the use of mouse preclinical data, sample size, data availability, and comparisons of sex, age, and disease stage. As with other disease preclinical models, the ADPKD mouse models do not perfectly recapitulate the human disease; attempted full *Pkd1* and *Pkd2* null mouse is embryonic lethal, which has led to models such as the ones used here that are expressed in the kidney only or as an inducible model to closer mimic adult onset ADPKD.(Traykova-Brauch et al., 2008; Williams et al., 2014) Additionally, while human *PKD1* variants are more prevalent in ADPKD patients, we focused here on *Pkd2* affected mouse models due to data availability. Future expansion of this method to *Pkd1* models as appropriate data becomes available would allow for further validation of our results. While the *Pkd2^fl^*^/fl^;*Pax8*rtTA;TetO-Cre mouse model may best recapitulate adult ADPKD, only one kidney RNA-seq data set was available for this model, and only two kidney RNA-seq data sets were available for *Pkhd1*-Cre; *Pkd2^fl^*^/fl^. As with other in-silico methods, these results should be further tested in-vitro and in-vivo to confirm efficacy. Lastly, we prioritized drugs with available drug target data, limiting this study to drugs with known targets and other annotations available in the Drug Repurposing Hub. Drugs were not considered if they were not available in the perturbation database (e.g., tolvaptan) or if they were not FDA-approved.

Future directions for this work include confirming the ability of these compounds to reverse ADPKD phenotypes in preclinical models and eventually, safety and efficacy for long-term treatment in patients. Additional drug repurposing approaches (such as combination therapy approaches) and a detailed assessment of how these drugs might differ across patients of different ages, sexes, and disease stages are also critical.(Fisher et al., 2022) However, this study provides a proof of concept on how transcriptomic data from ADPKD mouse models can be leveraged to identify drug repurposing candidates and novel drug targets.

## Conclusions

We provide prioritized drug repurposing candidates predicted to reverse the ADPKD molecular phenotype that we identified in-silico by transcriptomic signature reversion of publicly available pre-cystic *Pkd2* ADPKD mouse model RNA-seq data. We further prioritized these drugs by FDA status, pathway enrichment, and comparing their drug targets to the differentially expressed genes in additional cystic ADPKD mouse model RNA-seq profiles. We, therefore, determine the drug repurposing candidates that may be effective for early or preventative treatment, as well as cystic ADPKD, that can be further tested in in-vitro and in-vivo models.

## List of Abbreviations

(ADPKD): ADPKD
(PC1): polycystin-1
(PC2): and polycystin-2
(LINCS): LINCS
(cAMP): cAMP
(KO): KO
(LFC): LFC
(FEA): FEA
(GO): GO
(GO:BP),: GO biological process
(GO:CC),: GO cellular component
(GO:MF)): and GO molecular function
(GSEA): GSEA
(WTCS): WTCS
(ES): ES
(NCS): NCS
(DSEA): DSEA
(NEKs): NEKs
(RAAS): RAAS
(ESRD): ESRD
(AKI): AKI
(CKD): CKD
(NSCLC): NSCLC
(OCD): OCD
(SNRI): SNRI
(HMGCR): HMGCR
(CKD-aP): chronic kidney disease-associated pruritus
(CaM-MLCK): calmodulin-dependent myosin light chain kinase
(NF-κB): nuclear factor kappa-light-chain-enhancer of activated B cells
(PPARα): peroxisome proliferator-activated receptor α

## Co-author email addresses

Elizabeth J. Wilk

Timothy C. Howton

Jennifer L. Fisher

Vishal H. Oza

Ryan T. Brownlee

Kasi C. McPherson

Hannah L. Cleary

Bradley K. Yoder

James F. George

Michal Mrug

Brittany N. Lasseigne

## Declarations

### Ethics approval and consent to participate

Not applicable.

### Consent for publication

Not applicable.

### Availability of data and materials

All data is available from public repositories as stated in the text. All code needed to reproduce the analyses in this study and resulting Data Files are available at https://doi.org/10.5281/zenodo.7640442(Wilk et al., 2023)

## Funding

This work was supported in part by R03 (OD R03OD030604) (to BNL), the UAB Lasseigne Lab Start-Up funds (to BNL), UAB’s AMC21 Multi-PI 2021 (to BNL, BKY, JFG, and MM), the UAB Pilot Center for Precision Animal Modeling (C-PAM) (1U54OD030167) (to BNL and BKY), UAB KURE (NIH R25 DK 115353) in partnership with UAB Childhood Cystic Kidney Disease Core Center (UAB-CCKDCC) (NIH U54 DK126087) (supported RTB and HLC), Mentored Experiences in Research, Instruction, and Teaching (MERIT) Program (NIH K12 GM088010) (supported KCM), and National Institutes of Health (NIH)-funded PKD Research Resource Consortium (U54DK126087, U54DK128128 to MM), grants from the Office of Research and Development, Medical Research Service, Department of Veterans Affairs (1-I01-BX004232-01A2 to MM) and the Detraz Endowed Research Fund in Polycystic Kidney Disease (to MM). The funders had no role in the design of the study and collection, analysis, and interpretation of data and in writing the manuscript.

### Authors’ contributions

EJW, MM, and BNL conceptualized the project and EJW, TCH, and JLF contributed to the methodology used. Mouse RNA-seq data was curated by TCH, KCM, and HLC and preliminary analyses conducted by KCM and HLC. Gene expression heatmap and volcano plot visualizations were contributed by RTB. Original code by JLF was adapted for FDA approval testing. All other analyses were coded and performed by EJW. Code was reviewed and validated for reproducibility by TCH, JLF, and VHO. BNL provided supervision, project administration, and funding acquisition. EJW, TCH, and BNL contributed to writing the first draft. EJW, TCH, JLF, VHO, RTB, KCM, HLC, BKY, JFG, MM, and BNL reviewed and edited writing. All authors read and approved the final manuscript.

## Acknowledgments

The authors thank the Lasseigne Lab members Tabea Soelter, Jordan Whitlock, Amanda Clark, Anisha Haldar, Avery Williams, Emma Jones, Nathaniel DeVoss, and Victoria Flanary for feedback throughout this study.

## Additional Files

**Additional file 1:**
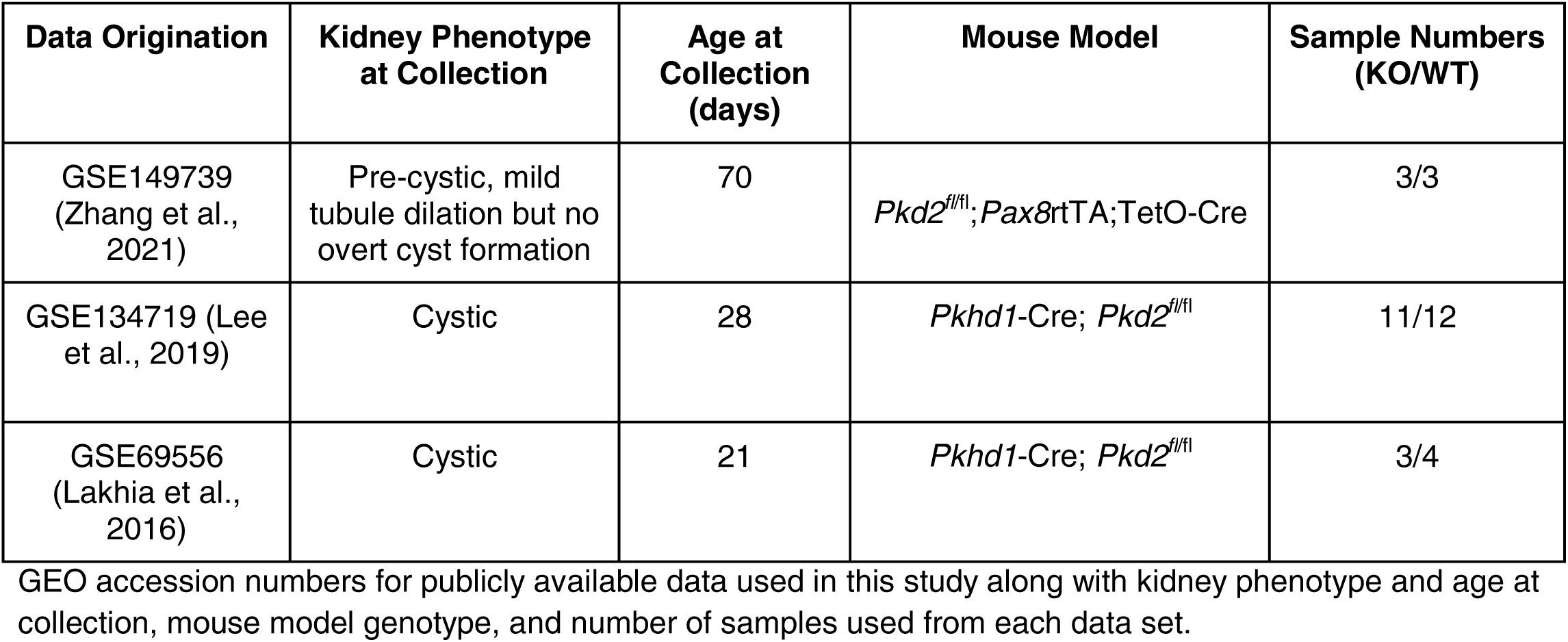
Summary of Data Sets used in this Study.

### Additional file 2: Enrichment of Unique Pre-Cystic DEGs and Overlapping Cystic Differentially Expressed Genes

**Additional file 2:**
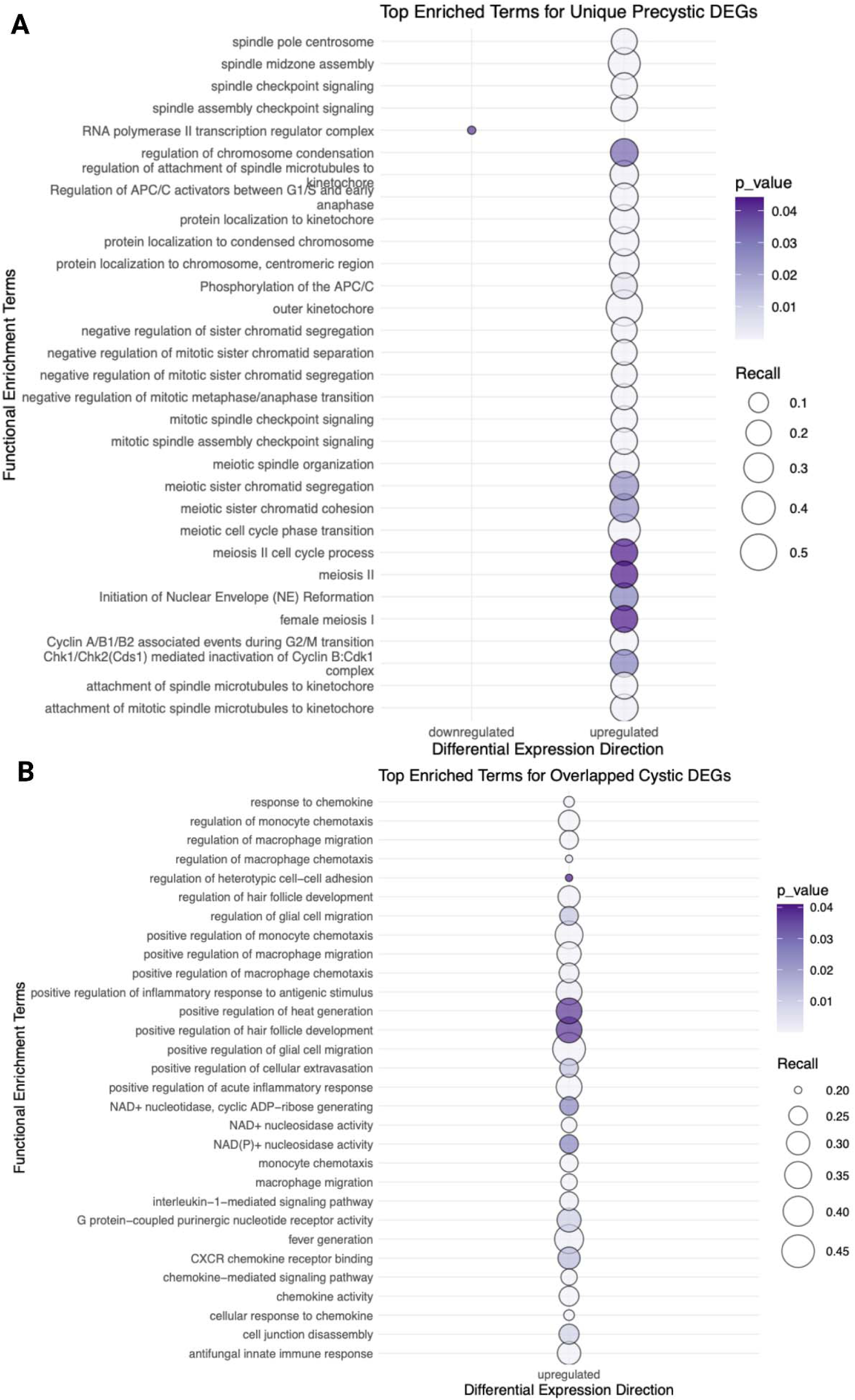
Pathway Enrichment of Pre-cystic (P70) vs Cystic (P21 and P28) Kidney Differential Gene Expression Bubbleplots showing enrichment for genes uniquely differentially expressed in the A) pre-cystic data set and B) both cystic data sets, where point size is the ratio of query genes to pathway gene set size.

### Additional file 3: Pathway Enrichment for Differentially Expressed Genes

**Additional file 3:**
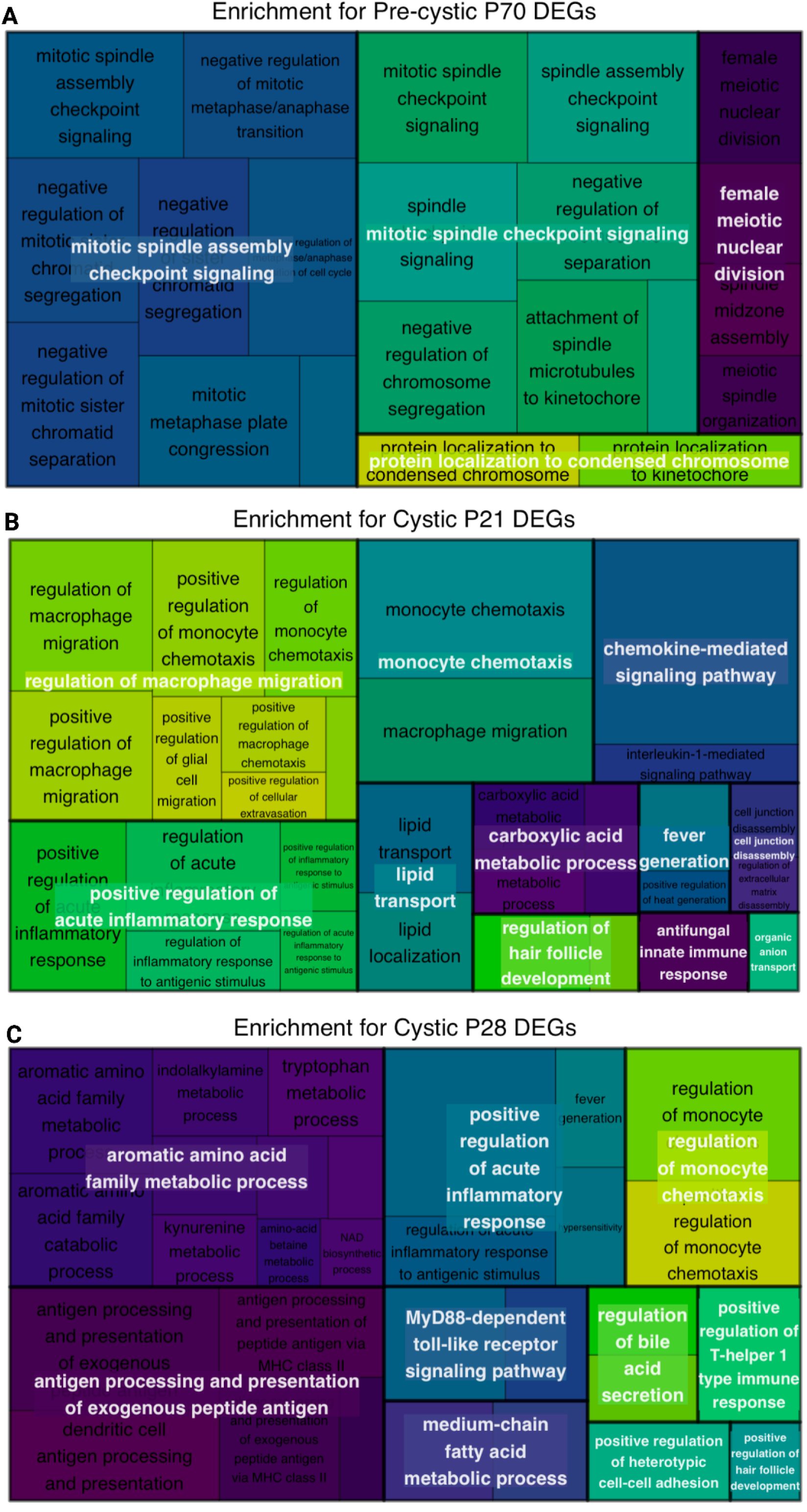
Differential Expression Analysis Treemaps of A) pre-cystic P70, B) cystic P21, and C) cystic P28 differentially expressed genes display enrichment nesting, with GO:BP parent terms consisting of largest boxes and enriched child terms reflecting enrichment term size nested within, for each signature.

### Additional file 4: Comparing Pre-Cystic and Cystic Transcriptomic Signatures

<u>Attached PDF</u>

**Additional file 4: Comparing Pre-Cystic and Cystic Transcriptomic Signatures** Results and discussion text comparing signatures derived from the differential expression of the pre-cystic and cystic mouse kidney data.

### Additional file 5: Transcriptomic Signatures for Pre-cystic and Cystic Disease Signature Reversion

**Additional file 5:**
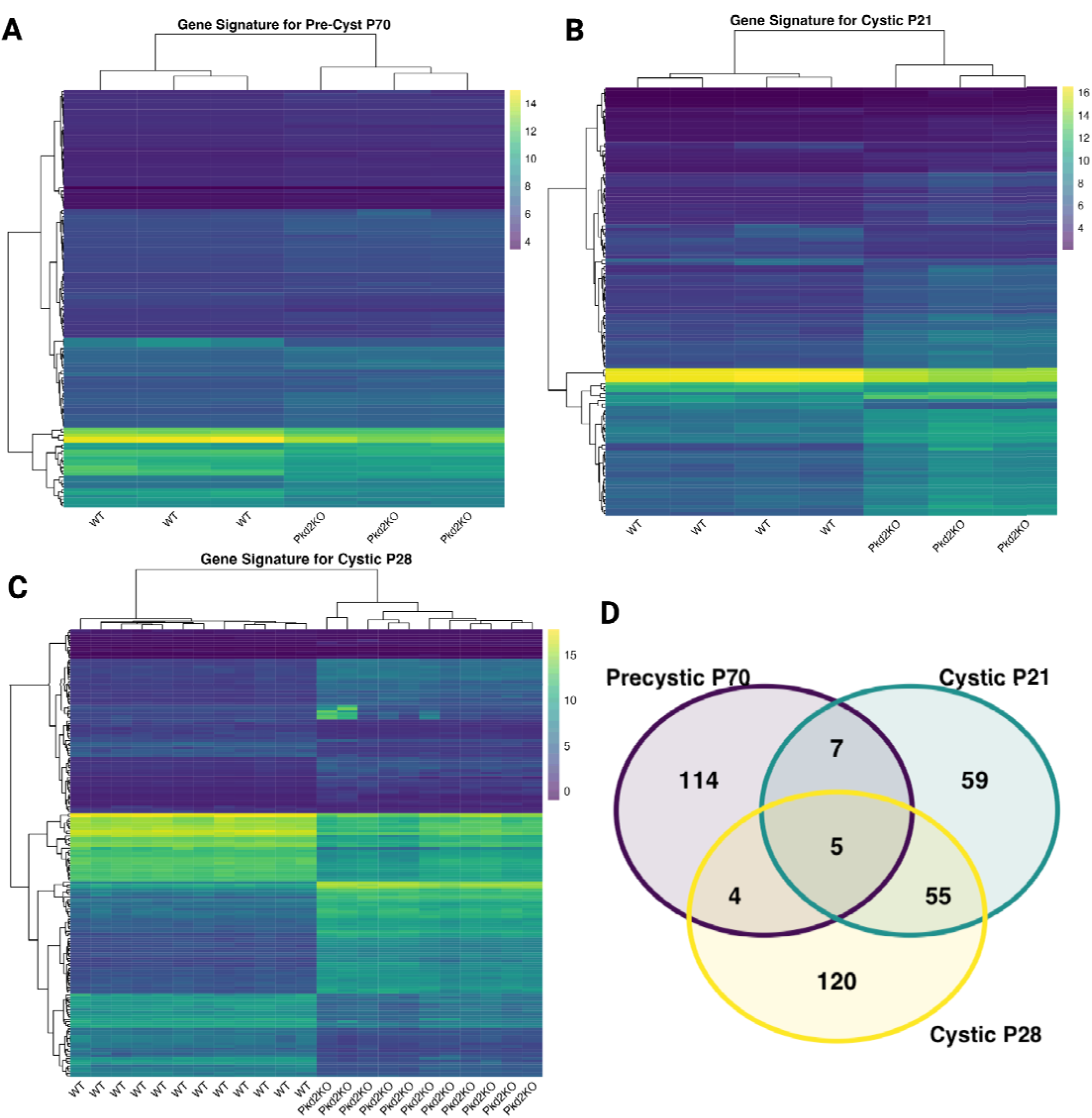
Transcriptomic Signatures for Pre-cystic and Cystic Disease Signature Reversion. Heatmaps of rlog gene expression counts for genes selected for transcriptomic reversion signatures (filtered by genes available in LINCS perturbation data and then top 100-200 absolute LFC genes) showing complete linkage hierarchical clustering of gene expression and samples for A) pre-cystic P70, B) cystic P21, and C) cystic P28. D) Venn diagram of transcriptomic reversion signatures from the 3 data sets showing overlap and unique genes for each signature.

