## Supplemental Information for "Prioritized polycystic kidney disease drug targets and repurposing candidates from pre-cystic and cystic mouse *Pkd2* model gene expression reversion"

Comparing Pre-Cystic and Cystic Transcriptomic Signatures

We then asked how the rlog-normalized gene counts for the selected genes in our disease gene signatures compared across samples in each data set. These disease signatures consisted of the 100-200 differentially expressed genes from each data set which mapped to LINCS measured or inferred genes as Entrez gene IDs (see Methods, Data files " deseq2_outputs").(Wilk et al., 2023) The expression of the signature genes for each data set for wild type and *Pkd2*-KO samples clearly clustered apart in each data set (Figure 2A-C).

We further compared the gene signatures from each data set to each other and found only 5 genes shared across the pre-cystic P70, cystic P21, and cystic P28 signatures (Figure 2D). Consistently upregulated genes in all 3 signatures included interleukin 1 receptor antagonist (*IL1RN*), serpin family A member 3 (*SERPINA3*), regulator of G protein signaling 16 (*RGS16*), and integrin subunit alpha M (*ITGAM*). The only consistently downregulated gene across all 3 signatures was parvalbumin, *PVALB,* which encodes a high-affinity calcium-binding protein found in the distal tubule.(Prot-Bertoye et al., 2022) We speculate that this downregulation is due to the decreased intracellular calcium availability resulting from polycystin dysfunction, as previously described (see Introduction).(Booij et al., 2017; Lemos & Ehrlich, 2018; Prot-Bertoye et al., 2022)

However, the two cystic transcriptomic signatures (P21 and P28) showed a high degree of similarity to each other, sharing 60 genes (47.6% and 32.2% of the P21 and P28 signatures in common with each other, respectively) (Figure 2D). Upregulated genes in both the P21 and P28 cystic signatures included 5 chemokine ligands/receptors *(CCR1, CCL8, CCL20, CXCL6,* and *CCN5*) and 5 interleukins (*IL1RN, IL6, LIF, IL18R1,* and *IL36A*). Chemokines and interleukins are commonly associated with PKD and, furthermore, as renal cysts progress in PKD and cystic vascularization increases, chemokine/immune responses increase and begin to reflect that of kidney injury.(Zimmerman et al., 2020) Additionally, cancer-associated genes *VGF* and *WNT7A*, and the forkhead box genes *FOXJ1* and *FOXN1* were also upregulated in both cystic data sets, reflective of the well-described overlaps between PKD pathogenesis and the hallmarks of cancer.(Seeger-Nukpezah et al., 2015) Additionally, it has been hypothesized that *PKD2* regulates *WNT7A* expression, and *Wnt7a* inhibition in *Pkd2*-/- cells has been observed to have decreased β-catenin levels.(Li et al., 2018)

The pre-cystic gene signature was much more unique, with 12 and 9 genes total shared with the cystic P21 and P28 signatures, respectively. Some of the upregulated genes that were found to be unique to the pre-cystic signature include cell division cycle genes *CDC25C* and *CDC20* and cyclin-family genes, including *CDK1* which was previously suggested as a driver of cyst proliferation in ADPKD.(Li et al., 2017; Zhang et al., 2021) Kinases unique to the upregulated pre-cystic signature included *PBK, BUB1, TTK, MELK, FGR,* and aurora kinases A and B (*AURKA* and *AURKB*). Aurora kinase A is particularly well-studied for its involvement in PKD and has been researched as a potential target for PKD treatment.(Booij et al., 2017; Plotnikova et al., 2011) Although chemokines were much more abundant in the cystic gene signatures (6 and 8 chemokine genes in P21 and P28 data sets, respectively, compared to 3 in the pre-cystic P70 signature)*, CCL18, CXCL10,* and *CCR5* were uniquely retained and specific to the pre-cystic signature after signature selection. Genes distinctly downregulated in the pre-cystic signature included DNA damage genes (*DDIT4* and *GADD45G*) and cytochrome P450 genes (*CYP2A13, CYP27B1,* and *CYP2A6*). Interestingly, *WNT11* was also downregulated only in the pre-cystic signature, and there is evidence that *WNT11* is critical for kidney tubules and renal function; C57Bl6 *Wnt11*−/− mice that survived to adulthood exhibited enlarged kidney tubules, altered kidney function, and prominent glomerular cysts.(Nagy et al., 2016)

<u>Discussion</u>

Additionally, we also found that *Wnt7a*, previously characterized for its role in WNT signaling in *Pkd2*-/- cells, was upregulated in both cystic datasets (P28 and P21).(Li et al., 2018) Foreseeably, there were more similarities in the differential expression of the P28 and P21 cystic data sets compared to the P70 pre-cystic data set (Figure 1A). All three datasets shared a total of 22 differentially expressed genes and the cystic data sets shared 387 differentially expressed genes. Consistently upregulated genes across all the data sets included many involved in immune pathways characteristic of ADPKD (*Ccl2, Ccl7, Ccr5, Cd44,* and the colony stimulating factor *Csf1*).(Vasileva et al., 2021) Our pathway enrichment analysis on all three data sets confirmed previously noted pathways involved in ADPKD pathogenesis, including inflammation, MAPK signaling, chemokine activity/receptor binding, and metabolic pathways.(Jouret & Devuyst, 2020; Podrini et al., 2020; Ye et al., 2016)
